# Representation learning of somatic missense mutation landscapes across cancer types

**DOI:** 10.64898/2026.10.25.761717

**Authors:** Zelin Wang, Minfang Cai, Zhewen Li, Zhenyu Lin, Yuanyuan Yin, Zhongyan Chen, Haiyan Yan, Zhen Quan, Jianyou Liao, Yiqing Zheng

**Affiliations:** Guangdong Provincial Key Laboratory of Cancer Pathogenesis and Precision Diagnosis and Treatment, Joint Big Data Laboratory, Department of Medical Oncology, Shenshan Medical Center, Memorial Hospital of Sun Yat-sen University, Shanwei, 516600, China; Medical Research Center, Sun Yat-sen Memorial Hospital, Sun Yat-sen University, Guangzhou, 510120, China; Department of Precision Medicine, Shenshan Medical Center, Sun Yat-sen Memorial Hospital, Sun Yat-sen University, Shanwei, 516600, China

## Abstract

Somatic missense mutations are pervasive in cancer genomes, yet existing studies have largely compressed them into mutation burden, mutation signatures, or driver gene annotations. Such representations obscure mutation-specific context and higher-order dependencies, leaving the latent biological structure of somatic mutation landscapes insufficiently characterized by systematic, data-driven approaches. Here we present MMRL, a pan-cancer **M**issense **M**utation **R**epresentation **L**earning framework that learns the representations directly from somatic missense mutation profiles. By integrating mutation-associated features including gene identity, trinucleotide context, AlphaMissense pathogenicity, amino acid substitution and protein positional information, MMRL uses self-supervised learning to capture higher-order patterns among missense mutations across ∼50,000 tumor samples from diverse cancer types. Across three pan-cancer cohorts, MMRL embeddings consistently achieved higher cross-validated prognostic discrimination than TMB and remained significantly associated with overall survival after adjustment for cancer type and TMB, as well as age where available. This prognostic value was also observed among patients treated with immune checkpoint inhibitors. Unsupervised clustering analysis of MMRL embeddings identified nine mutation subtypes in the TCGA pan-cancer cohort, with distinct survival outcomes, cancer-type compositions and mutational signature profiles, indicating that the learned embedding space captured clinically and biologically meaningful structure. Attention-based analyses further indicated that MMRL captured context-dependent mutation patterns beyond mutation frequency and predicted pathogenicity. Together, MMRL provides a data-driven strategy for decoding somatic missense mutation landscape and may facilitate precision oncology.

## Introduction

Cancer is driven by the accumulation of somatic genomic alterations^1^. Large-scale cancer sequencing projects have generated extensive catalogs of somatic mutations across diverse tumor types, providing an unprecedented opportunity to investigate how mutation patterns are associated with clinical phenotypes^2–6^. One of the most widely used measures derived from mutation data is tumor mutational burden (TMB), typically defined as the number of somatic non-synonymous mutations within a specified genomic region. TMB has been extensively evaluated as a predictive biomarker of response to immune checkpoint inhibitors across multiple cancer types^7^. However, as an aggregate measure of mutation burden, TMB does not account for relationships among individual mutations and therefore may not provide a sufficiently rich representation of a tumor’s mutational profile. This limitation highlights the need for approaches that can preserve mutation-level information while capturing complex mutation patterns in tumor.

Recent advances in representation learning have enabled complex biological entities, including genes, proteins, mutations, and patient samples, to be transformed into dense numerical embeddings that preserve latent biological relationships^8–10^. Compared with manually curated features, learned representations can capture nonlinear patterns and high-dimensional dependencies from large-scale data. In cancer genomics, recent studies have begun to learn mutation-level representations from somatic mutation profiles. Attention-based and multiple-instance learning approaches have encoded individual mutations and aggregated them into tumor-level representations for tasks such as tumor-type classification, tumor-subtype characterization and microsatellite-instability prediction^11,12^, while neural embedding models have also been developed to infer mutational signatures from targeted-panel data^13^. However, these approaches have largely been optimized for predefined supervised tasks. A generalizable framework that learns tumor representations directly from pan-cancer missense mutation profiles through self-supervised learning, and evaluates their utility for both survival prediction and subtype characterization, remains underexplored.

Here, we developed MMRL, a self-supervised pan-cancer framework that learns tumor-level representations directly from somatic missense mutation profiles. MMRL integrates gene identity, trinucleotide mutational context, AlphaMissense pathogenicity, amino acid substitution and relative protein position, and was trained on ∼50,000 tumor samples spanning diverse cancer types. We evaluated the learned embeddings across whole-exome and targeted-panel cohorts for survival prediction, prognostic stratification after immune checkpoint inhibitor treatment, tumor subtyping and biological interpretation. Our analyses showed that MMRL embeddings captured clinically and biologically meaningful variation beyond TMB, supporting a data-driven approach to tumor characterization from somatic missense mutation landscapes.

## Methods

### Data collection and processing

Cancer mutation data and matched clinical data were downloaded from cBioPortal^14^ (https://www.cbioportal.org/). Only tumor tissue samples were retained. Duplicate samples and germline variants were removed, and the analysis was restricted to missense mutations. Samples were eligible for representation learning if they contained at least 10 missense mutations affecting at least three distinct genes.

Somatic mutation data for TCGA were obtained from the Multi-Center Mutation Calling in Multiple Cancers (MC3) Public MAF file^15^ (mc3.v0.2.8.PUBLIC.maf.gz), and matched clinical annotations were obtained from the TCGA Clinical Data Resource^16^ (TCGA-CDR). AlphaMissense^17^ pathogenicity annotations were obtained from the publicly available AlphaMissense prediction dataset (https://console.cloud.google.com/storage/browser/dm_alphamissense). The cancer gene set was downloaded from the IntOGen database^18^, and the false driver gene set was obtained from Bailey et al^3^. TMB data were derived from Thorsson et al^2^.

For survival analyses among ICI-treated patients, progression-free survival was evaluated in 315 patients from five cohorts: mel_iatlas_liu_2019 (n=122)^19^, rcc_iatlas_immotion150_2018 (n = 106)^20^, paad_iatlas_prince_2022 (n=46)^21^, brca_iatlas_anders_2022 (n=25)^22^, ccrcc_iatlas_choueiri_2016 (n=16)^23^. Overall survival was evaluated in 491 patients from seven cohorts: blca_iatlas_imvigor210_2017 (n=242)^24^, mel_iatlas_liu_2019 (n=122)^19^, paad_iatlas_prince_2022 (n=46)^21^, brca_iatlas_anders_2022 (n=25)^22^, mel_iatlas_hugo_ucla_2016 (n=25)^25^, ccrcc_iatlas_choueiri_2016 (n=16)^23^, and gbm_columbia_2019 (n=15)^26^. Additional pan-cancer survival analyses were conducted in the msk_impact_2017^27^, tmb_mskcc_2018^28^ and msk_chord_2024^29^ cohorts.

### MMRL architecture

Each tumor sample was represented as an unordered set of missense mutations, truncated or padded to a fixed length of 255. Each mutation was characterized by gene symbol, trinucleotide context, AlphaMissense pathogenicity score, wild-type and mutant amino acids, and relative protein position. The gene vocabulary comprised curated cancer genes and the 8,000 most frequently mutated genes, with all other genes mapped to an <OTHER> token. Gene symbol and trinucleotide context were encoded using learnable embedding layers with dimensions of 256 and 128, respectively. The AlphaMissense pathogenicity score was projected into a 64-dimensional vector using a linear transformation. Wild-type and mutant amino acids were encoded as 64-dimensional vectors using a shared amino-acid embedding matrix. Relative protein position was projected into a 32-dimensional vector using a separate linear layer. The six resulting feature vectors were concatenated along the feature dimension to form a 608-dimensional mutation-level feature vector. This concatenated vector was subsequently passed through a fusion layer consisting of a linear projection, layer normalization, ReLU activation and dropout (p = 0.3), producing a unified 512-dimensional mutation vector. The fused vectors were then processed by a six-layer pre-norm transformer encoder with a hidden dimension of 512, eight attention heads, a feed-forward dimension of 1,024, GELU activation and a dropout rate of 0.3. Padding positions were excluded from self-attention using a sample-specific padding mask. The final mutation vectors were aggregated using pooling by multihead attention with eight learnable seed vectors. Each seed independently attended to the complete set of transformer outputs, producing eight 512-dimensional pooled vectors. These vectors were averaged to obtain the sample-level embedding:

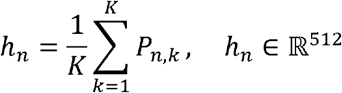

where *P _n, k_* denotes the vector generated by pooling seed k for tumor sample n. The resulting vector *h_n_* was used as the final mutation embedding for downstream analyses.

### Self-supervised learning

MMRL was optimized using two complementary self-supervised objectives: masked gene modeling and contrastive learning. For masked gene modeling, genes at a subset of mutation positions were replaced with a dedicated mask token, while the remaining features were retained. The model was trained to recover the original gene identities from the remaining mutation context using a cross-entropy loss calculated only at masked positions. Genes ranked among the 500 most frequently mutated genes were masked with a probability of 0.15, genes ranked from 501 to 3,000 were masked with a probability of 0.25, and less frequent genes were masked with a probability of 0.40. If no mutation was masked, one position was randomly selected to ensure at least one prediction target per sample. Let *Ω_n_ denote* the set of masked positions in tumor sample n. The masked gene modeling loss was defined as:

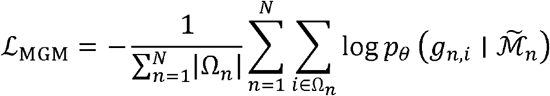

Where N is the batch size, *g_n,i_* denotes the original gene identity at masked position i in tumor sample n, *M̃_n_* is the corrupted mutation set and pθ is produced by a gene-classification head applied to the corresponding transformer output.

For contrastive learning, two independently augmented views were generated from each tumor sample. In each view, 10–30% of benign mutations were randomly removed, followed by independent frequency-aware gene masking. Each view was then processed by the shared encoder and attention-pooling module, and the resulting 512-dimensional vector was mapped to a 128-dimensional vector through a two-layer projection head with an ReLU activation. For each anchor, the other view of the same tumor served as the positive, whereas all views from others in the batch served as negatives. The normalized temperature-scaled cross-entropy loss was defined as:

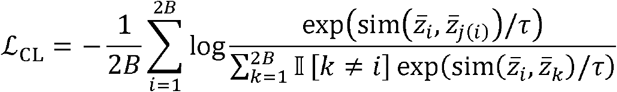

where *z̄_i_* denotes an *L_2_*normalized projected representation, *j(i)*identifies the other augmented view of the same tumor sample, and sim(.,.) denotes cosine similarity. The temperature parameter τ was set to 0.2.

The total self-supervised training objective was defined as a weighted combination of the masked gene modeling and contrastive learning losses:

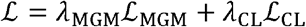

where the masked gene modeling and contrastive loss weights were set to 1.0 and 0.1, respectively. During training, the masked gene modeling loss was calculated from the first augmented view, whereas both augmented views were used to calculate the contrastive loss.

Models were trained using the AdamW optimizer with a learning rate of 1×10^-4^, a weight decay of 1×10^-4^, and a batch size of 256. Training was performed for up to 500 epochs using a 15% validation split. The checkpoint with the lowest validation masked gene modeling loss was retained, and training was stopped if the validation loss did not improve for 30 epochs.

### Survival analysis

Penalized Cox proportional-hazards models were fitted using the CoxPHFitter implementation in the lifelines python package. Penalizer values were set to 0.01 for the TMB-only and age-only models and to 0.10 for the MMRL-only and MMRL-containing models. Five-fold cross-validation was stratified by survival-event status, with shuffling enabled and a fixed random seed of 42. Predictive discrimination was quantified in the held-out samples using Harrell’s concordance index (C-index), and performance was summarized as the mean ± standard deviation across the five folds. For each patient, an out-of-fold MMRL risk score was obtained from the predicted partial hazard of the MMRL-only Cox model. Multivariable Cox models included the out-of-fold MMRL risk score, log-transformed TMB and age where available as continuous covariates, together with cancer type as a categorical covariate. The models were fitted with a penalizer of 0.05. For Kaplan–Meier analysis, patients were classified as high risk if their out-of-fold MMRL risk score was greater than the median and as low risk otherwise. Survival distributions were compared using the log-rank test. An additional univariable Cox proportional-hazards model with risk group as a binary predictor was used to estimate the hazard ratio and 95% confidence interval for the high-risk versus low-risk group.

### Consensus clustering analysis

The embeddings were standardized to zero mean and unit variance using the StandardScaler implementation in scikit-learn python package. Principal component analysis was subsequently applied to the standardized embeddings, and the first 50 principal components were retained for clustering. Consensus clustering was performed using the ConsensusClusterPlus R package^30^. Candidate cluster numbers from K = 2 to K = 20 were evaluated. For each K, 1,000 resampling iterations were performed, with 80% of the samples randomly selected in each iteration. K-means clustering with Euclidean distance was used as the base clustering algorithm. Cluster stability was primarily assessed by examining the consensus matrix heatmaps, the cumulative distribution function (CDF) of consensus values, and the relative change in the area under the CDF curve (delta area) across candidate K values. Stable clustering solutions were characterized by clear block patterns in the consensus matrix heatmaps, increasing concentration of consensus values toward 0 and 1 in the CDF plots, and a plateau in the delta area curve indicating limited stability gain with further increases in K. Based on these criteria, K = 9 was selected as the final clustering solution.

### Mutational signature analysis

PASS-filtered somatic single-nucleotide variants from the TCGA MC3 mutation file were classified into SBS96 categories using SigProfilerMatrixGenerator^31^ with the GRCh37 reference genome in exome mode. The SBS96 matrix was fitted to COSMIC SBS signatures (v3.5) using Analyzer.cosmic_fit in SigProfilerAssignment^32^ with exome renormalization enabled. The resulting activity matrix contained the estimated number of mutations assigned to each COSMIC SBS signature in each sample. Samples with a total assigned SBS activity of zero were excluded. For each retained sample, the activity assigned to each SBS signature was divided by the total activity across all SBS signatures to obtain the relative SBS exposure proportion. Mean relative SBS exposure proportions were calculated within each mutation subtype. Signatures detected in at least 2% of samples were ranked by their total assigned activity and mean relative exposure, and top 20 signatures were displayed. Subtype-associated SBS signatures were identified by comparing each subtype with all remaining subtypes using a one-sided Mann–Whitney U test. P values were adjusted using the Benjamini–Hochberg method, with FDR < 0.05 considered significant. Heatmap, dot plot and violin plot were generated using matplotlib python library.

### Attention-based analysis

Mutation-level attention weights were extracted from the pooling-by-multihead-attention (PMA) module. Attention weights were averaged across attention heads and pooling seed vectors, and attention was normalized within each sample to sum to one. normalized attention was summed to obtain gene attention scores. Subtype-specific gene attention was defined as the total gene attention scores within a subtype, normalized by the number of samples in that subtype. Genes mutated in at least five samples across the cohort were ranked within each subtype by subtype-specific gene attention, with mean attention among mutated samples and mutation frequency used as secondary ranking criteria. The top three genes per subtype were selected for visualization. In the cohort-wide analysis, mutation frequency was defined as the proportion of samples with at least one mutation in a given gene, and gene-level attention was defined as the mean gene attention among mutated samples. Gene-level AlphaMissense pathogenicity was calculated by first averaging mutation-level scores within each sample and then averaging across mutated samples. Associations of gene-level attention with mutation frequency and AlphaMissense pathogenicity were assessed using Pearson correlations.

## Results

### Overview of the MMRL framework

We developed MMRL, a pan-cancer missense mutation representation learning framework that encodes each tumor sample as a set of missense mutations (Fig 1A). MMRL integrates multiple mutation-level features, including gene symbol, trinucleotide context, AlphaMissense pathogenicity, wild-type and mutant amino acids, and relative protein position (Fig 1B). These features are embedded and fused into mutation-level vectors, which are processed by six transformer encoders to capture interactions among mutations within each sample. Pooling by multihead attention is then used to aggregate mutation-level vectors into a 512-dimensional embedding. MMRL was trained using two complementary self-supervised objectives. First, masked gene modeling required the model to predict randomly masked genes from the remaining mutation context, encouraging the encoder to learn mutation co-occurrence and contextual dependency patterns. Second, contrastive learning was used to enhance representation robustness by generating two augmented views of the same mutation set through random removal of a fraction of benign mutations. In total, MMRL was trained on ∼50,000 tumor samples spanning diverse cancer types and sequencing platforms, including targeted-panel, whole-exome and whole-genome sequencing.

**Fig 1.**
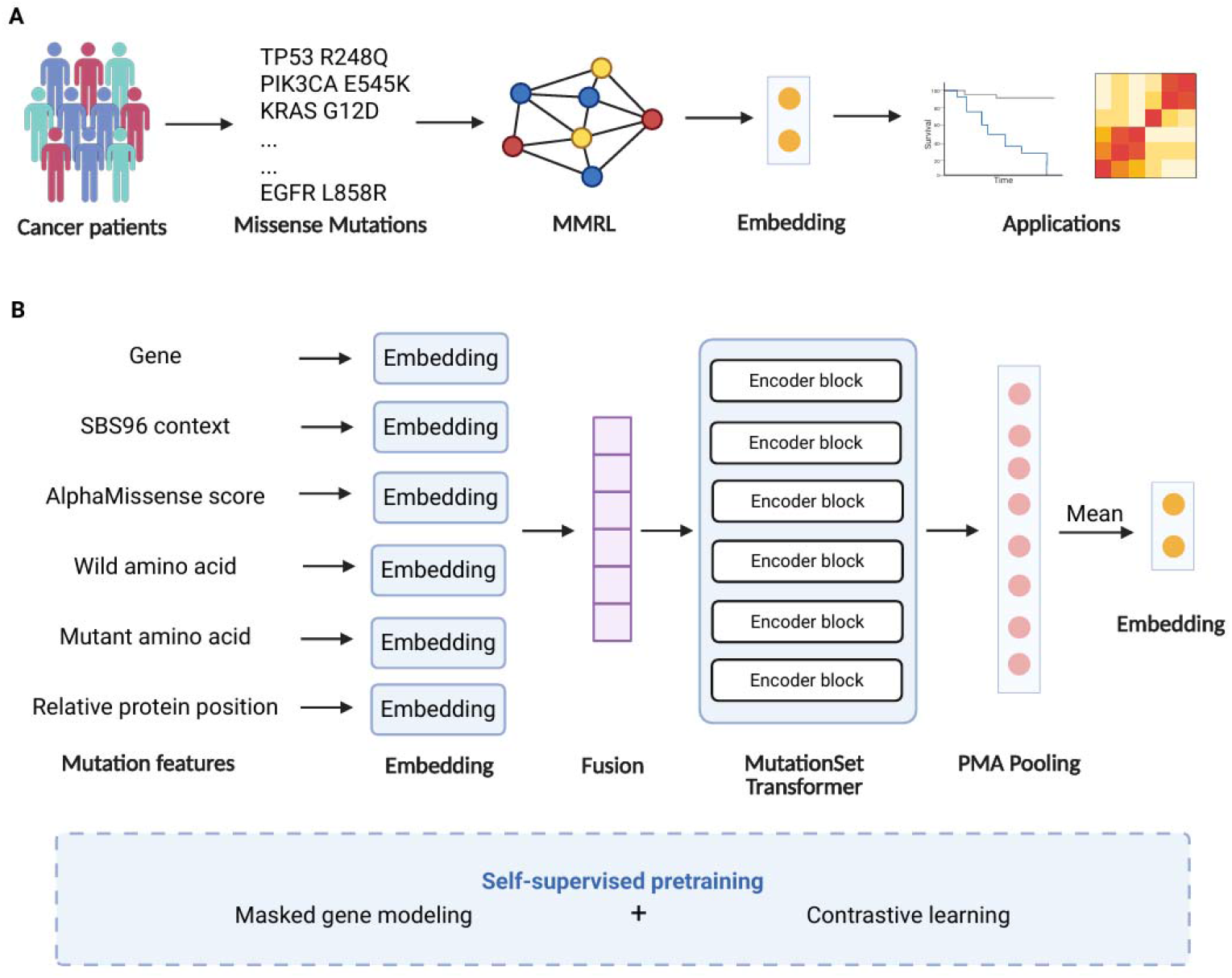
Overview of the MMRL framework. **(A)** MMRL learns representations from missense mutation set in tumor samples and generates embeddings for downstream applications. **(B)** Mutation features, including gene, SBS96 context, AlphaMissense score, wild-type and mutant amino acids, and relative protein position, are embedded and fused, then encoded by the MutationSet Transformer and aggregated by PMA pooling. The model is trained using masked gene modeling and contrastive learning.

### MMRL embeddings capture richer prognostic signals than TMB

To evaluate whether the learned missense mutation representations captured clinically relevant information, we first assessed their performance in pan-cancer survival prediction using TCGA samples. We found that TMB alone showed limited prognostic performance, with a C-index of 0.597 ± 0.013. Age achieved a higher C-index of 0.628 ± 0.010. In contrast, MMRL achieved a C-index of 0.657 ± 0.02, outperforming both TMB and age. The combination of TMB and MMRL yielded a slightly higher C-index of 0.658 ± 0.020. Further integration of the MMRL with age and TMB resulted in the best predictive performance, with a C-index of 0.674 ± 0.019 (Fig 2A). We next performed multivariable Cox regression to assess whether the MMRL-derived risk score was independently associated with overall survival after adjustment for cancer type, TMB and age. The results show that TMB was not significantly associated with overall survival (P = 0.14). In contrast, age remained strongly associated with survival risk (P = 7.50 × 10□^41^). Importantly, the MMRL was also significantly associated with overall survival (P = 3.20 × 10□^13^), with an increased hazard ratio per one standard deviation increase in the risk score (Fig 2B). We further used out-of-fold prediction scores to stratify TCGA pan-cancer patients into high-and low-risk groups. Patients assigned to the high-risk group had significantly worse overall survival than those in the low-risk group, with a hazard ratio of 2.50 (95% CI, 2.30–2.71) and a log-rank P value of 1.20 × 10□^111^(Fig 2C). Together, these results indicate that MMRL captures prognostic signals beyond those explained by age and TMB.

**Fig 2.**
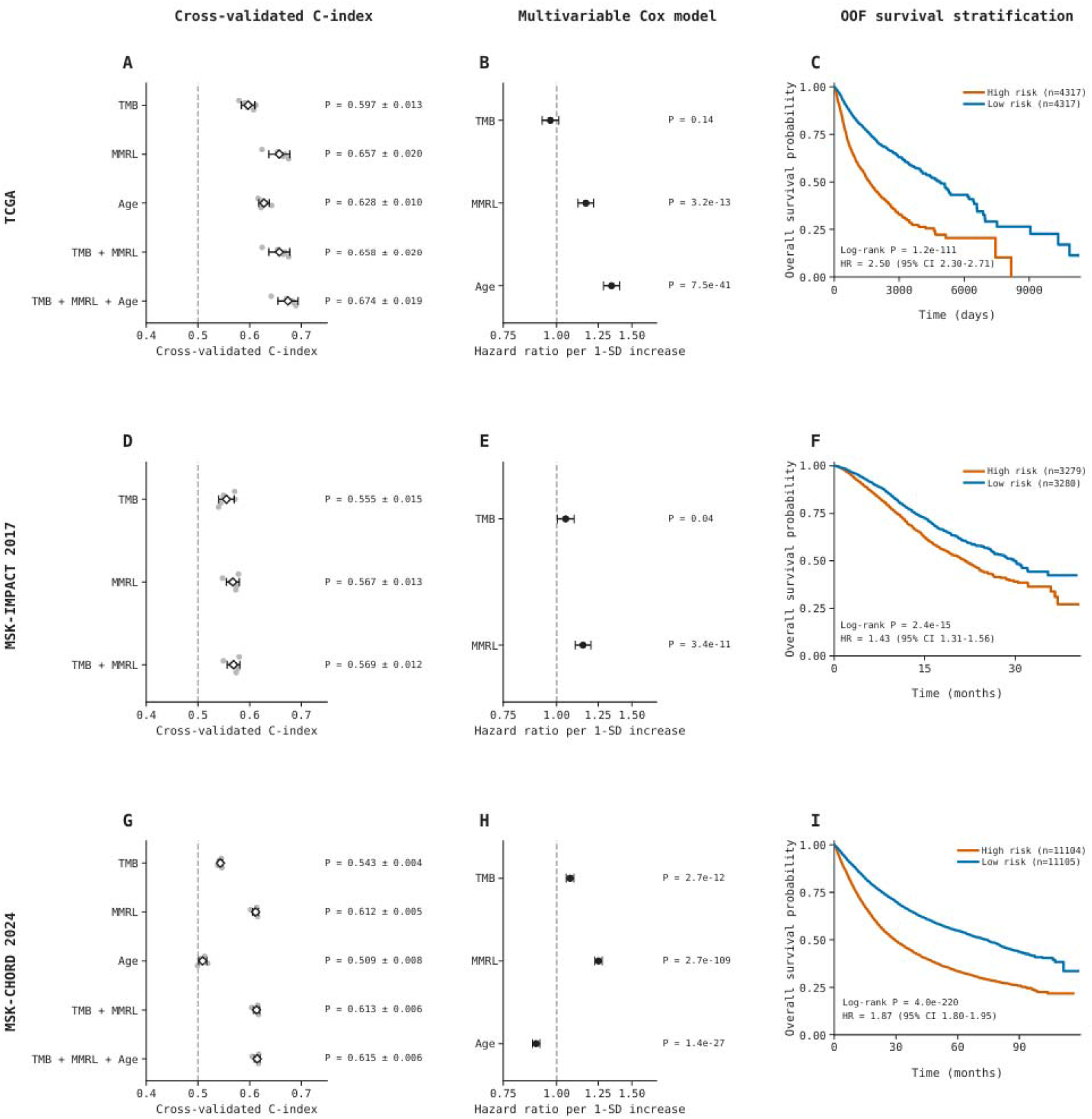
Prognostic performance of MMRL across multiple pan-cancer cohorts. **(A, D, G)** Cross-validated C-index (mean ± SD) of different models in the TCGA, MSK-IMPACT 2017, and MSK-CHORD 2024 cohorts. **(B, E, H)** Multivariable Cox regression showing hazard ratios (HRs) per 1-SD increase with 95% CIs. **(C, F, I)** Kaplan–Meier survival curves for OOF-defined high- and low-risk groups.

We also assessed whether this prognostic utility generalized to targeted-panel sequencing cohorts. In the MSK-IMPACT 2017 pan-cancer cohort, the TMB achieved a C-index of 0.555 ± 0.015, whereas the MMRL achieved a slightly higher C-index of 0.567 ± 0.013. Combining the MMRL with TMB slightly improved the C-index to 0.569 ± 0.012 (Fig 2D). Multivariable Cox regression analysis shows both TMB and MMRL were significantly associated with overall survival after adjustment of cancer type (TMB, P = 0.04; MMRL, P = 3.4 × 10□¹¹; Fig 2E). Out-of-fold risk stratification also separated patients into high- and low-risk groups with significantly different survival outcomes (HR = 1.43; 95% CI, 1.31–1.56; log-rank P = 2.4 × 10□¹□; Fig 2F). In the MSK-CHORD 2024 cohort, the TMB showed a C-index of 0.543 ± 0.004 and age showed a C-index of 0.509 ± 0.008, whereas MMRL achieved a substantially higher C-index of 0.612 ± 0.005. Combining MMRL with TMB resulted in a slightly improved C-index of 0.613 ± 0.006, and further incorporating age slightly improved the performance to 0.615 ± 0.006 (Fig 2G). In multivariable Cox analysis, the MMRL remained strongly associated with survival after adjustment for age and TMB as well as cancer type (P = 2.7 × 10□¹□□, Fig 2H). Consistently, out-of-fold MMRL-based risk stratification identified high- and low-risk groups with markedly different survival outcomes (HR = 1.87; 95% CI, 1.80–1.95; log-rank P = 4.0 × 10□²²□; Fig 2I). Overall, these findings demonstrate that MMRL retains prognostic utility across sequencing methods.

### MMRL improves prognostic stratification after *immune* checkpoint blockade

To evaluate whether MMRL embeddings were clinically informative in the immunotherapy prediction, we assessed their ability to predict survival after immune checkpoint inhibitor (ICI) treatment in three pan-cancer cohorts, including a pooled WES ICI cohort, MSKCC 2018, and MSK-CHORD 2024. In the pooled WES ICI cohort, MMRL showed greater discrimination for overall survival than TMB, with a C-index of 0.638 ± 0.021 compared with 0.539 ± 0.037 for TMB. Adding TMB to MMRL did not further improve performance (C-index, 0.637 ± 0.021; Fig. 3A). In multivariable Cox analysis, the MMRL remained significantly associated with post-ICI overall survival after adjustment of TMB, cancer type and studies (P = 5.0 × 10^-7^, Fig 3B). Consistently, out-of-fold risk scores separated patients into groups with markedly different outcomes: patients in the high-risk group had substantially worse survival than those in the low-risk group (HR, 2.06; 95% CI, 1.62–2.61; log-rank P = 2.2 × 10□□; Fig. 3C). For progression-free survival analysis, MMRL achieved a higher C-index than TMB (0.585 ± 0.029 versus 0.551 ± 0.033), whereas combining MMRL with TMB provided no additional improvement (0.583 ± 0.029; Fig. 3D). The MMRL remained associated with worse progression-free survival after adjustment of TMB, cancer type and studies (P = 0.02, Fig. 3E). Out-of-fold risk stratification further identified patients with distinct progression-free survival outcomes, with an HR of 1.67 for the high-risk group relative to the low-risk group (95% CI, 1.29–2.18; log-rank P = 9.8 × 10□□; Fig. 3F).

**Fig 3.**
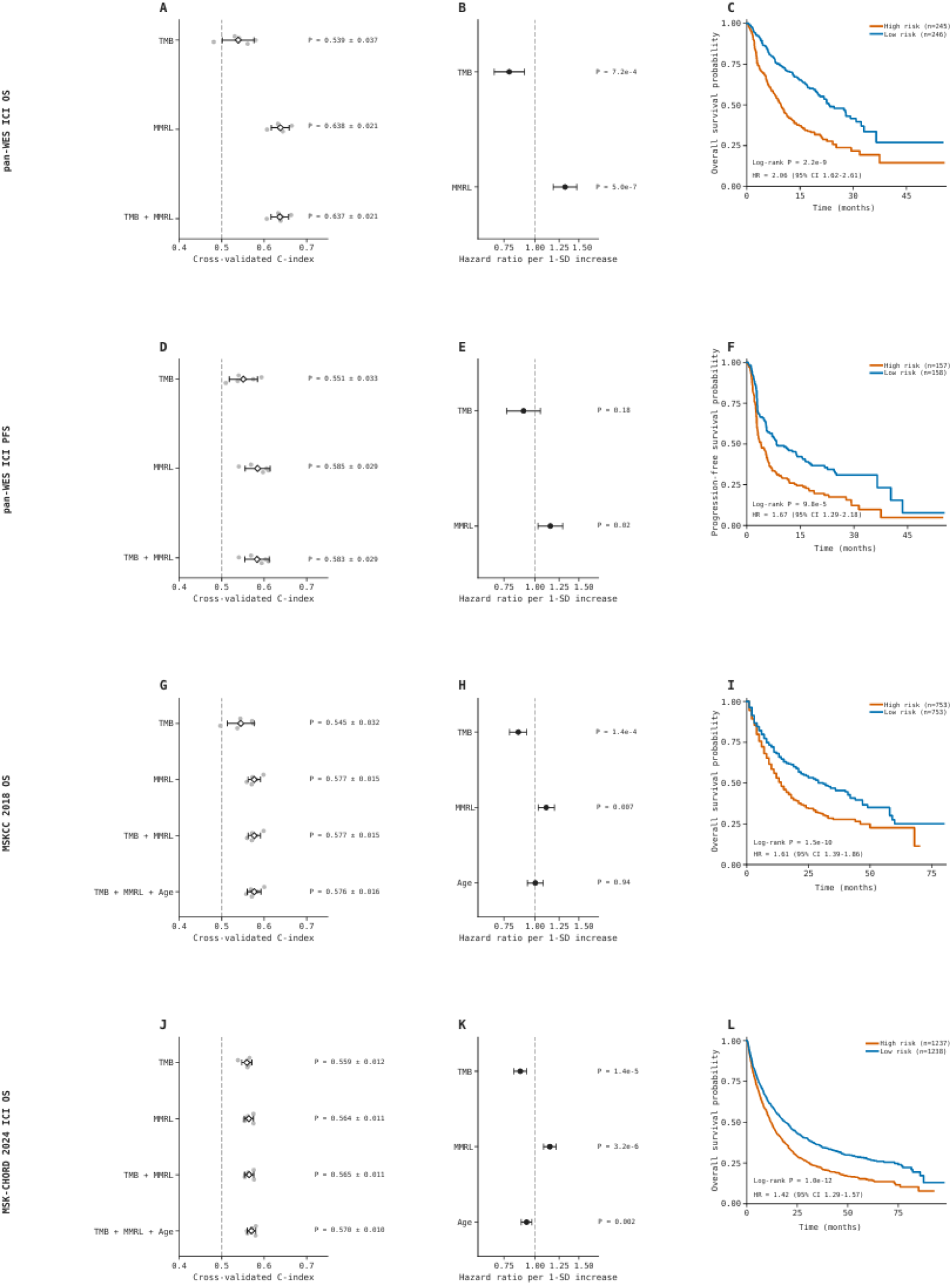
Prognostic performance of MMRL across ICI cohorts. **(A, D, G, J)** Cross-validated C-index (mean ± SD) of different models in the pan-WES, MSKCC 2018, and MSK-CHORD 2024 cohorts. **(B, E, H, K)** Multivariable Cox regression showing hazard ratios (HRs) per 1-SD increase with 95% CIs. **(C, F, I, L)** Kaplan–Meier survival curves for OOF-defined high- and low-risk groups.

The prognostic value of MMRL was reproduced in another two pan-cancer cohorts. In the MSKCC 2018 cohort, MMRL improved the cross-validated C-index from 0.545 ± 0.032 for TMB to 0.577 ± 0.015. Models combining MMRL with TMB, with or without age, showed similar discrimination (0.577 ± 0.015 and 0.576 ± 0.016, respectively; Fig. 3G). Both TMB and MMRL remained associated with overall survival in multivariable analysis, whereas age was not significant (Fig. 3H). Out-of-fold risk stratification separated patients into high- and low-risk groups with an HR of 1.61 (95% CI, 1.39–1.86; log-rank P = 1.5 × 10□¹□; Fig. 3I). In MSK-CHORD 2024, MMRL yielded a modest increase in discrimination over TMB alone (C-index, 0.564 ± 0.011 versus 0.559 ± 0.012). Combining MMRL with TMB resulted in a C-index of 0.565 ± 0.011, which increased to 0.570 ± 0.010 after the inclusion of age (Fig. 3J). The MMRL-derived risk score remained significantly associated with overall survival after adjustment of both TMB, age and cancer types (P=3.2 × 10□^6^, Fig. 3K). Patients classified as high risk again had worse survival than those classified as low risk (HR, 1.42; 95% CI, 1.29–1.57; log-rank P = 1.0 × 10□¹²; Fig. 3L). Together, these results indicate that MMRL can improve prognostic stratification after immune checkpoint blockade in pan-cancer cohorts.

### Unsupervised clustering of MMRL embeddings reveals pan-cancer mutation subtypes

To investigate mutation patterns captured through MMRL, we performed consensus clustering based on the MMRL embeddings in the TCGA pan-cancer cohort. The consensus matrix exhibited a clear block-diagonal structure, supporting a stable nine-cluster solution (C1–C9; Fig. 4A and Figure S1). Kaplan–Meier analysis revealed marked differences in overall survival among the nine subtypes (P=3.38 × 10□□², Fig. 4B). In particular, C4 showed the most favorable survival outcomes, whereas C5 and C6 exhibited comparatively poorer outcomes. We further investigated their cancer-type composition (Fig. 4C and Table S1), and found C4 was enriched for thyroid carcinoma, with thyroid carcinoma accounting for approximately 27% of the cluster. C5 was predominantly composed of lower-grade glioma samples, which represented approximately 41% of this subtype. C6 was strongly enriched for lung cancers, including lung squamous cell carcinoma and lung adenocarcinoma, which accounted for approximately 39% and 33% of the cluster, respectively. C8 was characterized by enrichment of skin cutaneous melanoma and bladder urothelial carcinoma, representing approximately 27% and 23% of the cluster. C9 contained a prominent mixture of gastrointestinal and endometrial tumors, including stomach adenocarcinoma, colon adenocarcinoma, and uterine corpus endometrial carcinoma, accounting for approximately 27%, 21%, and 18%, respectively. In contrast, C1, C2, C3, and C7 consisted of more heterogeneous combinations of cancer types, without a single overwhelmingly dominant lineage. Moreover, we also found C6, C8, and C9 showed relatively high median log-transformed TMB values, whereas C4 exhibited the lowest TMB and C5 showed a comparatively low-to-intermediate level (Fig 4D). Notably, clusters with similar TMB levels still differed in cancer composition and survival, suggesting that the MMRL embeddings encode not only the overall quantity of somatic mutations but also qualitative differences in mutation composition. We also found that C4 and C5 containing relatively younger patients, whereas C6 and C9 tended to include older individuals (Fig 4E). Clinical stage composition appeared to vary across subtypes, although interpretation was limited by the high proportion of samples with unknown stage (Fig. 4F). Sex distributions were generally comparable across most subtypes (Fig. 4G), although C6 showed a modest male predominance, consistent with its enrichment for lung cancers. Together, these results demonstrate that MMRL can capture biologically and clinically meaningful pan-cancer mutation patterns.

**Fig. 4.**
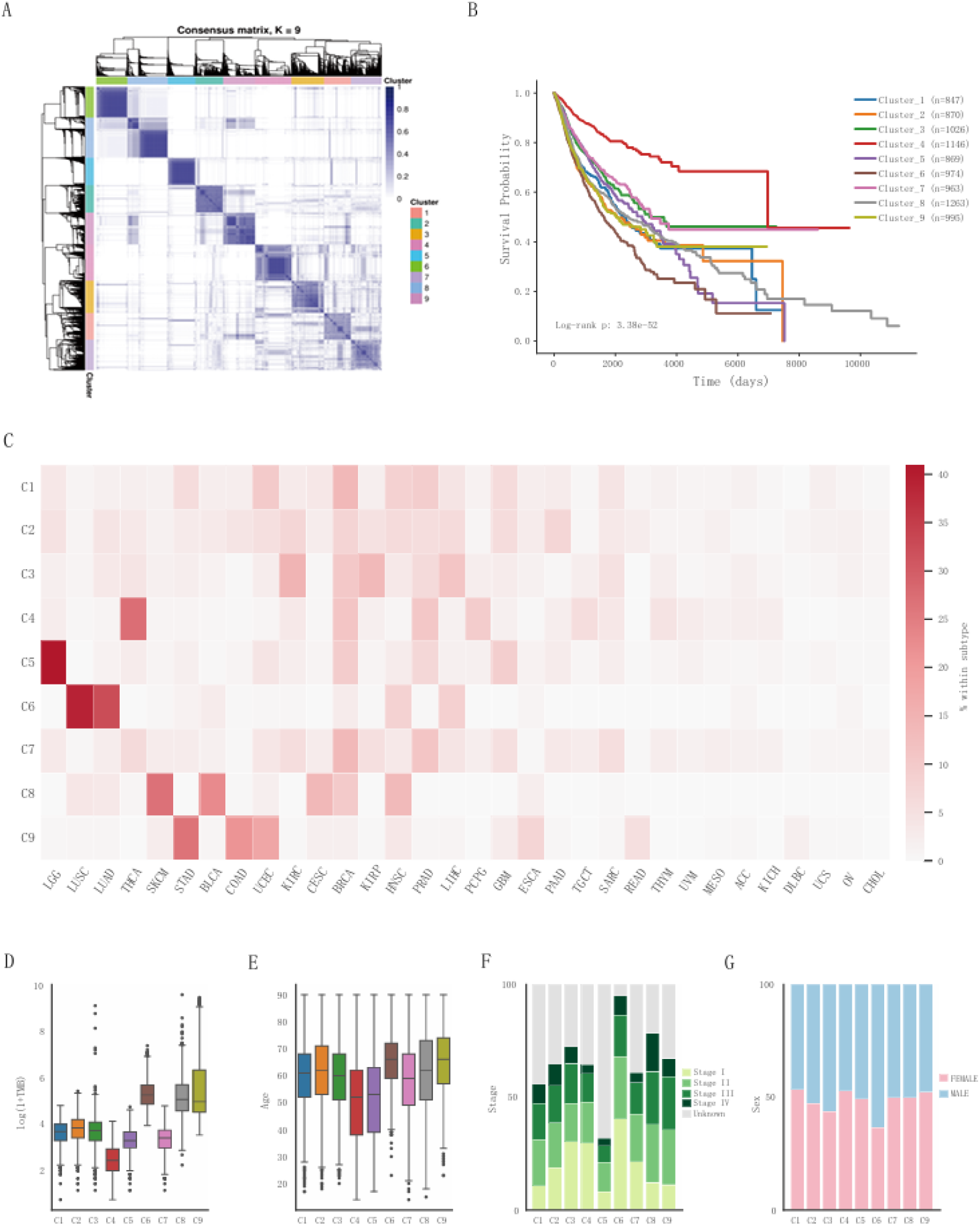
Pan-cancer mutation subtypes identified from MMRL embeddings and their clinical characteristics. **(A)** Consensus clustering matrix showing the nine-cluster solution (C1–C9). **(B)** Kaplan–Meier curves comparing overall survival among the nine subtypes; *P* value was calculated using the log-rank test. **(C)** Heatmap showing the cancer-type composition of each subtype. **(D–G)** Distribution of TMB, age, clinical stage, and sex across the nine subtypes, respectively.

### MMRL-defined subtypes exhibit distinct mutational signatures

We next examined SBS mutational signatures to characterize the mutational processes associated with the MMRL-defined subtypes. The SBS exposure landscape differed markedly across subtypes (Fig. 5A), indicating substantial heterogeneity in the mutational processes represented within the MMRL embedding space. Differential exposure analysis further revealed both subtype-specific and shared signatures (Fig. 5B). For example, C6 was significantly enriched for the tobacco smoking-associated signature SBS4, consistent with its predominance of lung cancers, whereas C8 showed enrichment of UV-associated SBS7a and APOBEC-associated SBS13, concordant with its enrichment for melanoma and bladder cancer. In contrast, several subtypes shared broader clock-like signatures, with SBS1 enriched in C1, C2, C5 and C9, whereas SBS5 in C1, C2, C3, C7 and C9. These associations were also evident at the individual-sample level, with the top enriched signature for each subtype showing higher exposure in the corresponding group than in the remaining subtypes (Fig. 5C). Together, these findings show that MMRL-defined subtypes are distinguished by biologically interpretable mutational processes, indicating that the learned embedding space captures meaningful variation in tumor mutation profiles.

**Fig. 5.**
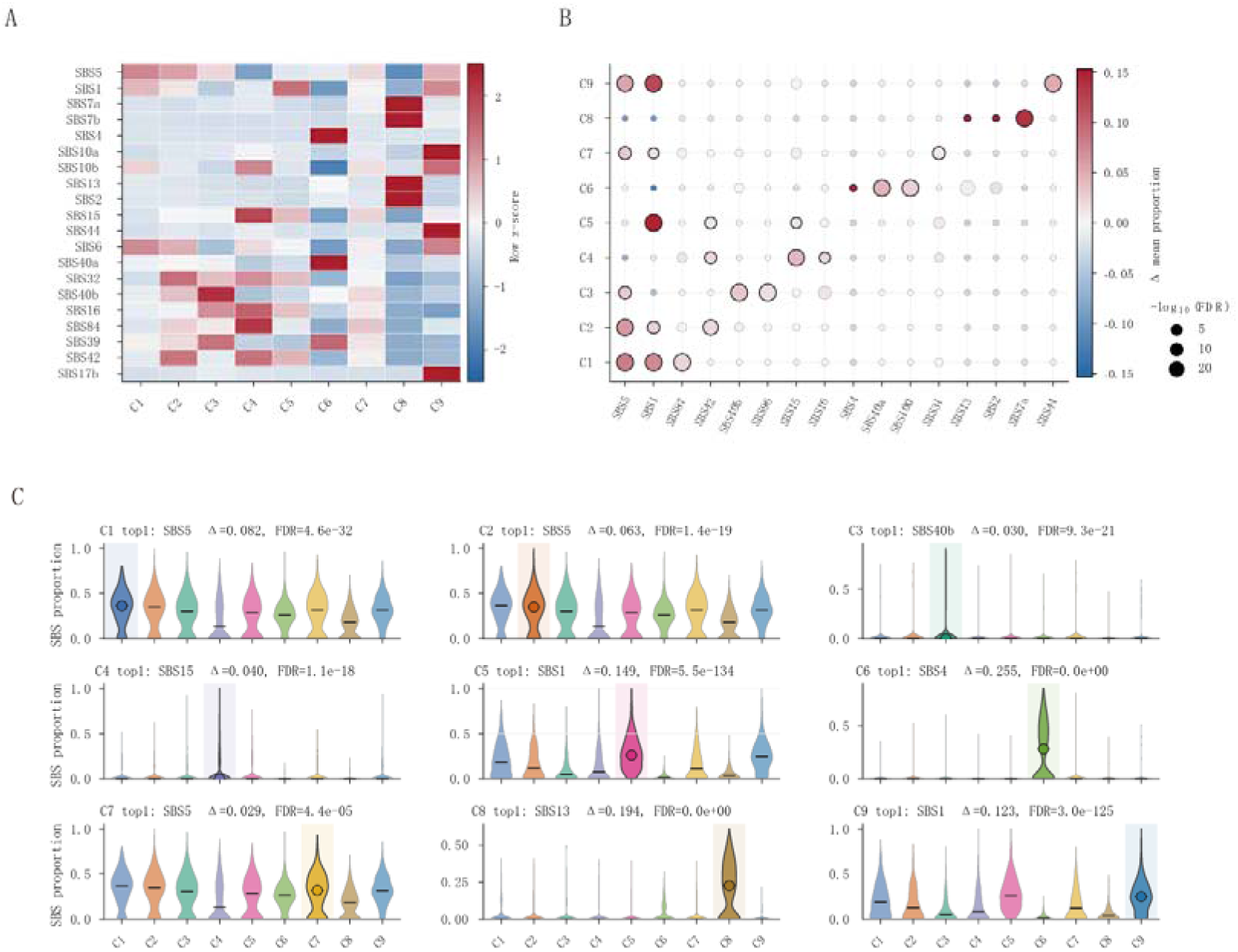
SBS mutational signature profiles of MMRL-defined subtypes. **(A)** Heatmap showing the relative exposure patterns of SBS mutational signatures across subtypes. Values represent row-wise *z*-scored mean SBS exposure proportions, with red and blue indicating relatively higher and lower exposure, respectively. **(B)** Bubble plot showing significant associations between SBS signatures and subtypes. Bubble color represents the difference in mean SBS exposure proportion between the indicated subtype and the remaining samples (Δ mean proportion), with red indicating enrichment and blue indicating depletion. Bubble size represents the significance of the association, expressed as −log□□(FDR). **(C)** Sample-level distributions of the top one SBS signature for each subtype.

### Attention analysis reveals subtype-dependent mutation patterns

To investigate the mutation patterns associated with the MMRL-defined subtypes, we aggregated mutation-level attention weights at the gene level and compared attention profiles across subtypes (Fig. 6A). The results showed that canonical cancer genes such as TP53, KRAS and EGFR received high attention across multiple subtypes, whereas some other genes were preferentially highlighted in specific subtypes. For example, C1 was distinguished by attention to HCFC1, DPF2 and ARHGAP4 in addition to TP53, whereas C7 was characterized by CLIC6 and SMARCA4 alongside TP53. Notably, C6 and C8 showed distributed attention profiles despite their strong enrichment for specific cancer types; a similar pattern was also observed in the more cancer-type-heterogeneous C9. These observations indicate that subtype-associated attention profiles were not simply determined by tumor lineage or canonical cancer genes. We next asked whether gene-level attention primarily reflected mutation frequency or AlphaMissense pathogenicity. Attention showed negligible correlation with mutation frequency (Pearson r = 0.01, Fig. 6B) and only a modest positive correlation with mean AlphaMissense pathogenicity (Pearson r = 0.25, Fig. 6C). These results suggest that MMRL prioritized mutations in a context-dependent manner, rather than simply ranking them according to recurrence or predicted pathogenicity.

**Fig. 6.**
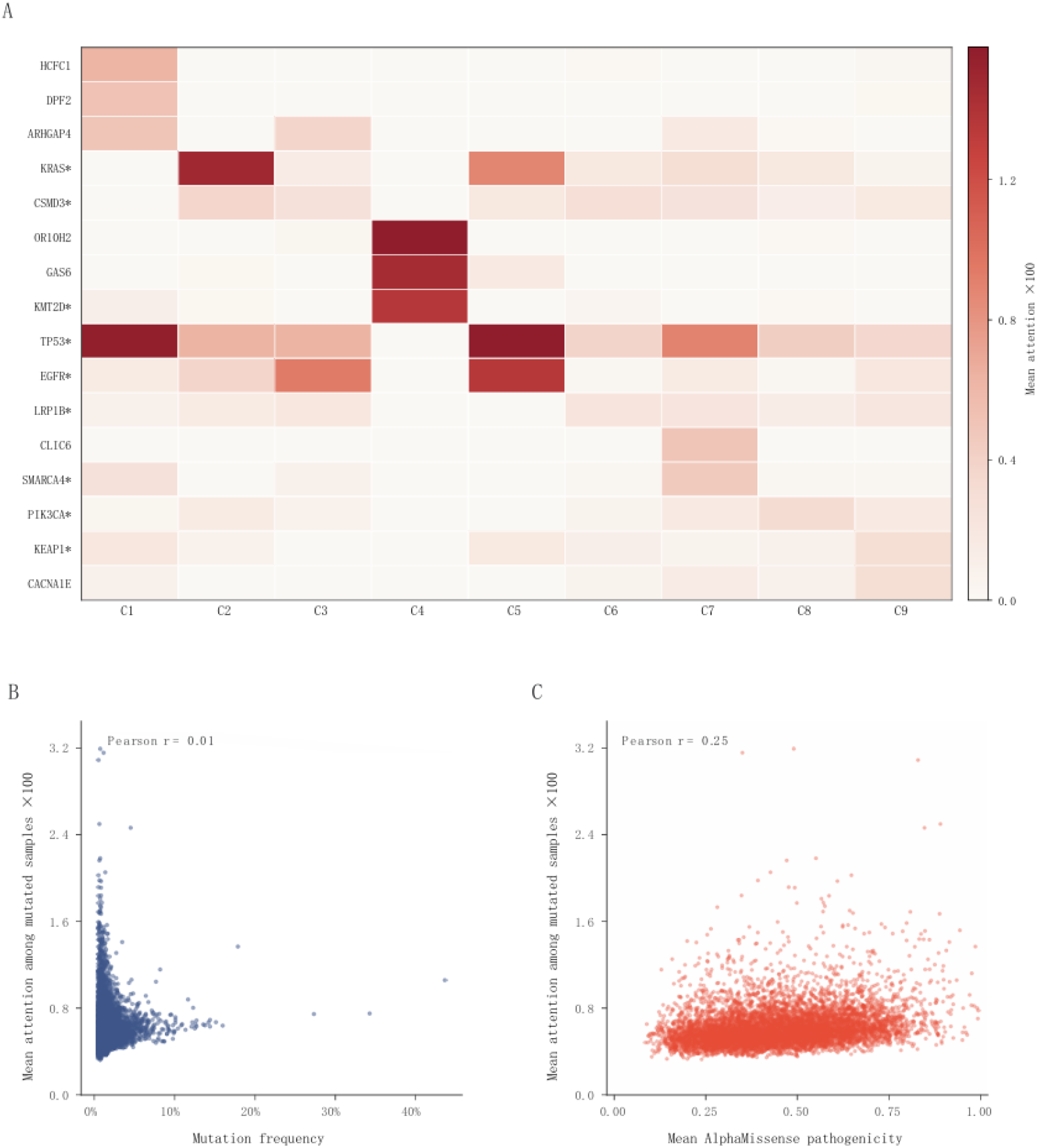
Subtype-specific gene attention patterns learned by MMRL. **(A)** Heatmap of attention-prioritized genes across the nine MMRL-defined subtypes. Color indicates subtype-level gene attention; asterisks denote known cancer genes. **(B)** Relationship between gene-level attention and mutation frequency across the full cohort. **(C)** Relationship between gene-level attention and mean AlphaMissense pathogenicity. Each point in (B) and (C) represents one gene.

## Discussion

Cancer somatic mutation profiles are commonly characterized using mutation burden^7,28,33,34^, recurrently altered genes^3,35–38^, or mutational signatures^6,39–41^. Although these approaches have provided important biological and clinical insights, each capture only part of a highly heterogeneous mutation landscape. For example, TMB captures the overall quantity of mutations but not their composition, whereas mutational signatures infer underlying mutational processes but lose gene- and protein-level context. Here, we developed MMRL to learn the representations of missense mutation profile of tumor sample by integrating gene identity, trinucleotide context, AlphaMissense pathogenicity, amino acid substitution and protein positional information.

Across three pan-cancer cohorts, MMRL consistently outperformed TMB in C-indices and remained significantly associated with overall survival after adjustment for TMB, cancer type and age where available. The minimal improvement observed when TMB was added to MMRL suggests that the learned representation contains prognostic information beyond that captured by TMB alone. One possible explanation is that, because MMRL is learned directly from tumor-specific missense mutation profiles through self-supervised learning, it may implicitly encode information related to the overall number of mutations, and therefore already incorporates part of the signal represented by TMB. At the same time, its superior prognostic performance suggests that MMRL may capture not only TMB but also higher-order features of the mutational landscape, such as the functional contexts, and co-occurrence patterns of missense mutations. By contrast, TMB compresses genomic complexity into a single quantitative measure, and its clinical interpretation is influenced by tumor type^28^, patient population^42^, assay methodology^43,44^, and treatment context^45^, which may partly account for its weaker and less transferable prognostic performance^46^. Importantly, MMRL also retained prognostic value among patients treated with immune checkpoint inhibitors, suggesting it may capture more granular information that is relevant to immunotherapy outcomes.

The unsupervised clustering and annotation of the MMRL embedding provided evidence that the learned representation captured biologically meaningful structure in the missense mutational landscape. The resulting nine subtypes differed in survival outcomes, clinical characteristics, and cancer-type composition, indicating that proximity within the embedding space was associated with clinically relevant tumor properties. The enrichment of several subtypes for specific cancer lineages further suggests that MMRL retained lineage-related mutational features. However, the presence of subtypes containing tumors from diverse anatomical origins indicates that the representation was not determined solely by tissue of origin and may also capture shared mutational patterns across conventionally distinct cancer types. SBS signature analysis further suggested that MMRL embeddings also retained information about underlying mutational processes. Moreover, the attention analysis demonstrates that MMRL captures mutation information beyond simple recurrence and AlphaMissense pathogenicity. Together, these findings support MMRL as a robust and biologically informative framework for learning the representations from missense mutation profiles.

This study has several limitations. First, MMRL was trained on retrospective cohorts with heterogeneous sequencing platforms, clinical annotations and treatment contexts. Although this heterogeneity supports broad applicability, it also introduces potential confounding by cohort, cancer type and assay design. Second, targeted-panel data are enriched for cancer-related genes, which may influence the learned gene representations. Finally, MMRL currently focuses on missense mutations and does not explicitly model other alteration classes, such as truncating mutations, copy-number alterations and structural variants. Future work could extend this framework such as integrating additional genomic alteration types may allow MMRL-like models to capture a more complete representation of tumor genomic alterations. In addition, experimental and functional analyses of attention signals may help determine whether model-prioritized missense mutations reflect previously unrecognized biological dependencies.

In summary, MMRL demonstrates that missense mutation sets encode richer information than TMB alone. By learning missense mutation landscape, MMRL connects mutational processes, prognosis, mutation subtypes and immunotherapy outcomes across cancer types. These findings support a shift from count-based mutation summaries toward representation-based interpretation of cancer mutation profiles.

## Author contributions

Y.Z. and J.L. conceived and supervised the project. Z.W. designed and developed the method, performed the analyses, and wrote the original draft. M.C., Z.L., Z.L., Z.C., Y.Y., H.Y. and Z.Q. contributed to the study design, data analyses and validation. All authors contributed to the review and editing of the manuscript and approved the final version.

## Data availability

The data analyzed in this study are publicly available through cBioPortal for Cancer Genomics (https://www.cbioportal.org/). The specific cancer cohorts and datasets used in the analyses are described in the Methods section.

## Code availability

The source code for MMRL, including model training, inference and saved model weights, is publicly available at https://github.com/charlin90/MMRL.

## Ethics approval and consent to participate

All data used are publicly available.

## Consent for publication

Not applicable.

