## Supplementary figures and images for "Representation learning of somatic missense mutation landscapes across cancer types"

### Figure S1.pdf

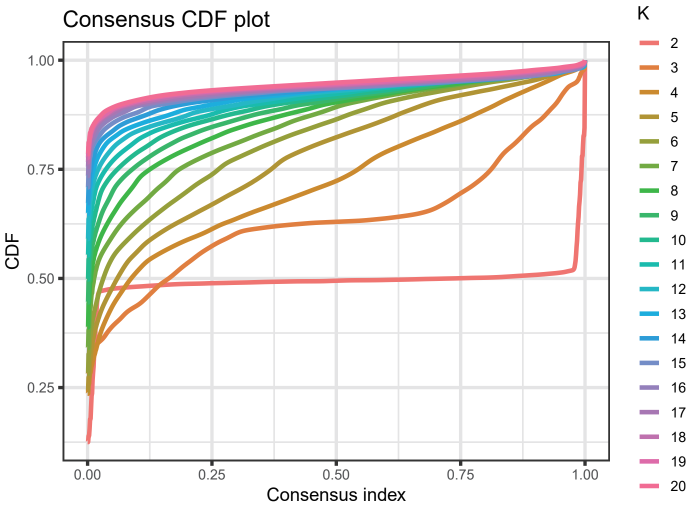

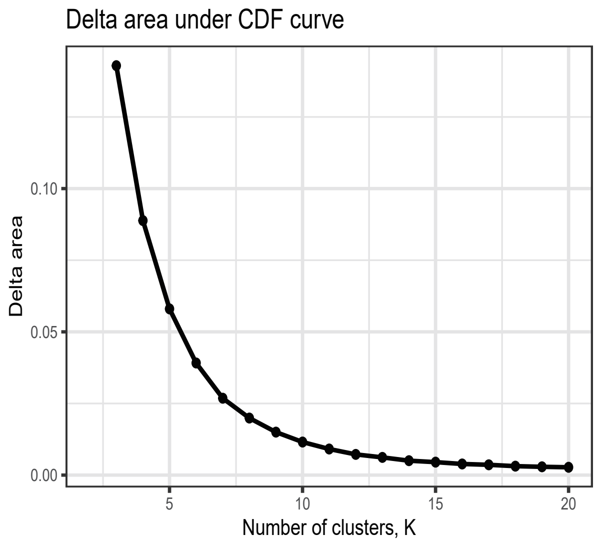
